# Divergent effects of C4a, C4a^desArg^, and thrombin on platelet aggregation and phosphorylation of ERK and Akt in human endothelial cells

**DOI:** 10.1101/2024.03.13.584877

**Authors:** Mengyao Liu, Vy K. Tran Luu, Hongbin Wang

**Affiliations:** Master Program of Pharmaceutical Sciences College of Graduate Study; California Northstate University, Elk Grove, CA 95757; Department of Pharmaceutical and Biomedical Sciences College of Pharmacy; California Northstate University, Elk Grove, CA 95757; Department of Basic Science College of Medicine; California Northstate University, Elk Grove, CA 95757

**Keywords:** Complement, C4a, C4a^desArg^, Thrombin, Extracellular signal-regulated kinases (ERK), Protein kinase B (Akt), Calcium influx, Platelet aggregation, Protease-activated receptor (PAR) 1 and 4

## Abstract

Prior studies have established C4a as an untethered ligand for protease-activated receptors (PAR)1 and PAR4, which can increase ERK phosphorylation and [Ca^2+^]_i_ influx in human endothelial cells (ECs). C4a^desArg^ is a stable metabolite produced from C4a through cleavage of an arginine at the carboxyl terminus by plasma carboxypeptidases B/N. PAR1 and PAR4 are typical receptors for thrombin and transduce cellular responses to the serine protease generated by the activation of coagulation pathways. Here, we aim to address whether C4a^desArg^ can induce the same effects as C4a through PAR1 and PAR4, and whether C4a and C4a^desArg^ can activate the same downstream signaling effectors as thrombin through PAR1 and PAR4.

We demonstrated that C4a^desArg^ induces ERK phosphorylation and [Ca^2+^]_i_ influx with the reduced efficacy as compared to C4a in human ECs. Distinct from C4a, C4a^desArg^-induced ERK phosphorylation was only inhibited by the PAR4 antagonist tcY-NH_2_, indicating that C4a^desArg^-mediated ERK phosphorylation is PAR4-dependent. Both C4a and C4a^desArg^ at a concentration of 3 μM failed to induce platelet aggregation. Moreover, both C4a and C4a^desArg^ induce significant Akt phosphorylation, whereas thrombin causes Akt dephosphorylation in human ECs.

Our study revealed that the absence of the C-terminal arginine in C4a decreases its efficacy and changes its preference for receptor of ERK and Akt activations in human ECs, suggesting that the C-terminal arginine of C4a might govern its binding specificity and/or affinity to PAR1 and/or PAR4. Unlike thrombin, both C4a and C4a^desArg^ fail to induce platelet aggregation at supraphysiological concentrations. Contrary to thrombin, both C4a and C4^adesArg^ induce significant Akt phosphorylation, indicating a unique role of C4a and C4^adesArg^ in inflammation and coagulation through their association with PAR1 and/or PAR4.

## Introduction

Complement C4 is a key component in the classical and lectin complement pathways, crucial for defending against microorganism infection, as well as clearing immune complexes and apoptotic cellular debris^(1, 2)^. Upon the activation of the classical or lectin complement pathways, activated C1s or MASP-1/2 from the respective classical or lectin pathways cleave the amino terminal part of the α-chain at a single site of complement C4 to generate C4a (77 aa) and C4b^(2)^. C4a was thought as the third anaphylatoxin although it is still in dispute^(3, 4)^. C4a has been reported to inhibit C3a-induced O^2-^ generation in guinea pig macrophages. Recombinant C4a has been shown to inhibit C3a- or C5a-induced chemoattractant and secretagogue functions in mast cells; inhibit C5a-induced neointima formation; and protect against hyperoxic lung injury ^(5–9)^. Our G protein-coupled receptor (GPCR) screening assay led to the identification of PAR1 and PAR4 as the receptors for C4a ^(10)^. Activation of PAR1 and PAR4 by C4a causes [Ca^2+^]_i_ influx, ERK phosphorylation, and the disruption of endothelial barrier function, consistent with a previous reports of intradermal injection of C4a inducing edema in human skin^(3,10)^.

C4a, a 77 aa peptide, is released into the bloodstream upon the activation of the classical or lectin complement pathway, lasting for a short period before being rapidly cleaved by plasma carboxypeptidases N /B (CPN/B) to its more stable des-arginated metabolite C4a^desArg^ ^(11)^. We posit that C4a^desArg^ may exert its pathophysiological functions more systemically than C4a, which may function only at the site of complement activation. It has been reported that C4a^desArg^ levels are significantly high in patients with IgA nephropathy and idiopathic pulmonary arterial hypertension (IPAH), serving as a plasma biomarker for complement activation in these disorders ^(12, 13)^. However, the exact role of C4a^desArg^ under physiopathological conditions has not yet been defined. In the present study, we aim to examine the effect of C4a^desArg^ on human platelet aggregation, ERK and Akt phosphorylation, and [Ca^2+^]_i_ influx in human ECs.

PAR1 and PAR4 belong to a subfamily of GPCRs, including PAR1, 2, 3, and 4 ^(14)^. PAR1 and PAR4 are typical receptors for thrombin, transducing cellular responses to the serine protease generated by the activation of coagulation pathways. Thrombin is locally generated at the sites of vascular injury, where it initiates responses in various cell types including platelets, endothelial, and smooth muscle cells. Thrombin is the key effector protease of the coagulation pathway, binding directly to PAR1 and PAR4 and exerting proteolytic cleavage of the N-terminal regions of the receptors. The newly formed N-terminus serves as a tethered ligand for the respective receptor, binding intramolecularly to the receptor’s second extracellular loop, causing a conformational change of the receptor that facilitates heterotrimeric G protein coupling and downstream signaling, leading to platelet activation and leukocyte-mediated EC reactions ^(15, 16)^. A synthetic six-amino acid peptide mimic of a PAR’s N-terminal tethered ligand is sufficient to activate the receptor and induce cellular responses similar to those generated from proteolytic cleavage of the receptor ^(17)^.

Our previous GPCR screening assay established that C4a^desArg^, like C4a, can act as an untethered ligand for PAR1 and PAR4, but with the reduced efficacy and potency^(10)^. Nonetheless, the effects of C4a^desArg^ on platelet aggregation and EC functions are still obscure. Our present data show that C4a^desArg^ can cause ERK phosphorylation and [Ca^2+^]_i_ influx in human ECs at concentrations as low as ∼10 nM. C4a^desArg^-induced ERK phosphorylation in human ECs could be inhibited by the PAR4 antagonist tcY-NH_2_, but not by the PAR1 antagonist RWJ56110, which is distinct from C4a-induced ERK phosphorylation that could be inhibited by both PAR1 and PAR4 antagonists.

To determine whether C4a/C4a^desArg^ and thrombin induce the same activation pattern of the downstream effector ERK and Akt, we compared Akt phosphorylation (p-Ser473) induced by C4a, C4a^desArg^, and thrombin. C4a and C4a^desArg^ time-dependently (5-30 min) and significantly induce Akt phosphorylation. In contrast, thrombin at concentrations from 0.1 to 1 nM (5-30 min) could significantly decrease Akt phosphorylation (dephosphorylation). The data suggest that C4a and C4a^desArg^ act differently from thrombin on PAR1/4 and their downstream signaling effectors. The study also revealed that C4a-induced phosphorylation of Akt and ERK is PAR1- and PAR4-dependent, whereas C4a^desArg^-induced phosphorylation of Akt and ERK is PAR4-dependent.

Taken together, our present studies confirm that C4a^desArg^ could induce ERK and Akt phosphorylation and [Ca^2+^]_i_ influx with the reduced efficacy as compared to C4a, indicating that the C-terminal arginine in C4a plays a role in defining the affinity and specificity to its receptor PAR1 and/or PAR4. In addition, C4a/C4a^desArg^-mediated Akt phosphorylation is distinct from thrombin in human ECs, suggesting that C4a/C4a^desArg^ and thrombin act differently on PAR1 or PAR4 and induce distinctive downstream effector functions despite acting on the same receptors PAR1 and/or PAR4.

## Materials and Methods

### Reagents

C4a (Catalog no. A106, Lot #16) and C4a^desArg^ (Catalog no. A107, Lot #10) were purchased from Complement Technologies (Tyler, TX). Antibodies for p44/42 MAPK (ERK1/2) (Catalog no. 9102), Akt (Catalog no. 9272), phosphor-p44/42 mitogen-activated protein kinase (MAPK) (extracellular signal-regulated kinase [ERK] 1/2) (Thr202/Tyr204) antibody (Catalog no. 4370), and phosphor-Akt (Ser473) (Catalog no. 9271) were purchased from Cell Signaling Technology, Inc. (Danvers, MA). PAR1 antagonist RWJ65110 (Catalog no. 2614) and PAR4 antagonist tcY-NH_2_ (Catalog no. 1488) were purchased from Tocris Bioscience (Minneapolis, MN). PAR1 antagonist SCH79797 (Catalog no. 1939) and PAR4 antagonist ML354 (Catalog no. 1439) were purchased from Millipore Sigma. Antibody for β-actin (Catalog no. A2228) and human α-thrombin (Catalog no. T6884-100N) was purchased from Sigma-Aldrich. All other reagents were bought from Fisher-Scientific except where noted.

### Cell culture

Human microvascular endothelial cell line of dermal origin (HMEC-1) was obtained from ATCC and cultured in growth medium consisting of MCDB 131 (Gibco) supplemented with 10% FBS (Sigma), 1 μg/mL hydrocortisone, 100 units/mL penicillin, 100 mg/mL streptomycin, 2 mM glutamine (Gibco), and 10 ng/mL epidermal growth factor (Fisher Scientific). The human hybrid cell line EA.hy 926 (ATCC), made via fusion of HUVEC with A549 lung adenocarcinoma cells, was maintained in DMEM with 10% conditioned FBS, penicillin (100 units/mL), streptomycin (100 mg/mL), and L-glutamine (2 mM).

### Preparation of platelet rich plasma (PRP) and platelet poor plasma (PPP)

With approved IRB protocol (approval # 818267) by the University of Pennsylvania Institutional Review Board (IRB), human blood was collected by vein puncture into tri-sodium citrate (6 volume + 1 volume acid citrate dextrose (ACD). Final pH = 6.5 and citrate concentration = 22 mM. PRP was obtained by centrifugation of whole blood (200 g, 20 min, 25 °C) and a platelet count was performed to confirm normal platelet numbers (2-4 × 10^8^ platelets/ml). PPP (platelet-poor plasma) was obtained by collecting supernatant from fraction of further centrifugation of PRP (1100 g, 5 min, 25 °C).

### Platelet aggregation assay

Platelet aggregation was measured using CHRONO-LOG® Model 700 Whole Blood/Optical Lumi-Aggregometers (Havertown, PA). The pre-warmed curvets were added with 250 μl of PRP with the speed of a stirrer at 1200 rpm at 37°C. PPP was used as a 100% aggregation control. The platelet aggregation was measured as % of transparency of PPP control.

### Western blots to determine the activations of extracellular signal-regulated kinase (ERK) and protein kinase B (Akt) in human ECs

HMEC-1 or EA.hy 926 cells were seeded into 24-well plates (200,000 cells per well) and cultured at 37 °C and 5% CO_2_ for ∼48 h until confluent. Following exchange of the growth medium with HBSS containing 2% BSA (cell culture grade), the cells were cultured for 16 h, and then treated with various concentrations of C4a, C4a^desArg^, or thrombin for the time indicated in the experiments, washed with cold DPBS, and lysed in 1× protein loading buffer. The samples were boiled for 5 min, separated by 10% SDS/PAGE, and analyzed by Western blot using antibodies for β-actin, phospho-extracellular signal-regulated kinase 1/2 (p-ERK1/2), phospho-Akt (Ser473), total ERK, or total Akt to detect p-ERK1/2, total ERK, p-Akt (Ser473), total-Akt or β-actin. Goat anti-mouse or goat anti-Rabbit HRP-conjugated secondary antibodies were used. In some experiments, IRDye^®^ 680RD Goat anti-Mouse IgG (H+L) or IRDye® 800RD Goat anti-Rabbit IgG (H+L) secondary antibodies were used. Western blot images were developed by enhanced chemiluminescence or by Odyssey DLx image system (LICOR). Immunoblots were quantified by densitometry using ImageJ software (NIH). ERK and Akt activation were expressed as the ratio of the densitometry of p-ERK/the densitometry of the total ERK or β-actin and the densitometry of p-Akt (Ser473)/the densitometry of total Akt or β-actin, respectively.

### Measurement of intracellular calcium concentration [Ca^2+^]_i_

Intracellular calcium concentrations measurement using a Fluo-4 NW calcium assay kit (Molecular Probes, Catalog no. F36206) was described elsewhere ^(10)^. All procedures were performed according to the manufacturer’s protocol. In brief, HMEC-1 cells were seeded into black-walled 96-well plates (40,000 cells per well) and cultured at 37 °C and 5% CO_2_ for ∼48 h until confluent. The growth medium was subsequently removed from the adherent cell cultures. After 100 μL of 1× dye loading solution (Fluo-4 NW with 2.5 mM probenecid) was added and the samples were incubated for 30 min at 37 °C, [Ca^2+^]_i_ was measured by fluorescence spectrometry using a Victor Spectrophotometer X4 with an excitation (Ex) at 485 nm/emission (Em) at 535 nm. In each assay, vehicle alone was used as a negative control and ionomycin (1.3 μM) as a positive control for each experiment to validate the experiment. The baseline relative fluorescence units (RFU_0_) for each well were recorded for 10 s. After addition of the test reagents, the RFU in each well was recorded every 2 s for 90 s. The RFU of each well at each time point was calculated as ΔRFU_t_ = RFU_t_ − RFU_0_. At each time point, the value of test sample RFU was obtained by subtracting the control vehicle RFU: Δ(ΔRFU)_t_ = test sample ΔRFU_t_ − Vehicle ΔRFU_t_. The peak value of Δ(ΔRFU)_t_ for each test sample was used to represent [Ca^2+^]_i_ influx of the test reagent.

### Statistical analysis

Data are expressed as mean ± SE. Statistical analyses were conducted using un-paired two-sided Student’s *t* test.

## Results

### C4a and C4a^desArg^ fail to induce platelet aggregation

Our prior screening assay for GPCR activation (DiscoverX) leads to identification of PAR1 and PAR4 as receptors for C4a and C4a^desArg^ ^(10)^. Thrombin has been well documented to activate PAR1 and PAR4 by generating newly exposed tethered ligands and cause platelet activation and aggregation. Synthetic six-amino acid peptides mimic of PAR1 or PAR4 N-terminal tethered ligand is sufficient to activate the receptors and induce cellular responses similar to the tethered ligand ^(17)^. Right after uncovering PAR1 and PAR4 as the receptors for C4a, we set to test whether C4a and C4a^desArg^ could trigger platelet aggregation through binding to PAR1 and/or PAR4. Our data revealed that both C4a and C4a^desArg^ at supraphysiological concentration of 3 μM were unable to induce human platelet aggregation (Fig. 1A and 1B), which was in line with a recent report by Han *et al*. ^(18)^. After adding C4a (3 μM) or thrombin (0.1 nM) to the platelet rich plasma (PRP) preparation, the platelet aggregation curve shows a negative “bump”, which indicates morphological changes of the platelets (Fig. 1A and 1C). C4a (3 μM) could induce substantial morphological changes of platelets, which resembles the effects induced by thrombin (0.1 nM). However, these morphological changes caused by C4a didn’t proceed to induce platelet aggregation. Thrombin at the concentration of 0.3 nM, 0.5 nM, and 1 nM induces ∼10% (Fig. 1D), ∼40% (Fig. 1E), and ∼100% (Fig. 1F) of platelet aggregation, respectively. Our study confirmed that C4a (3 μM) and C4a^desArg^ (3 μM) were unable to induce platelet aggregation.

**Figure 1.**
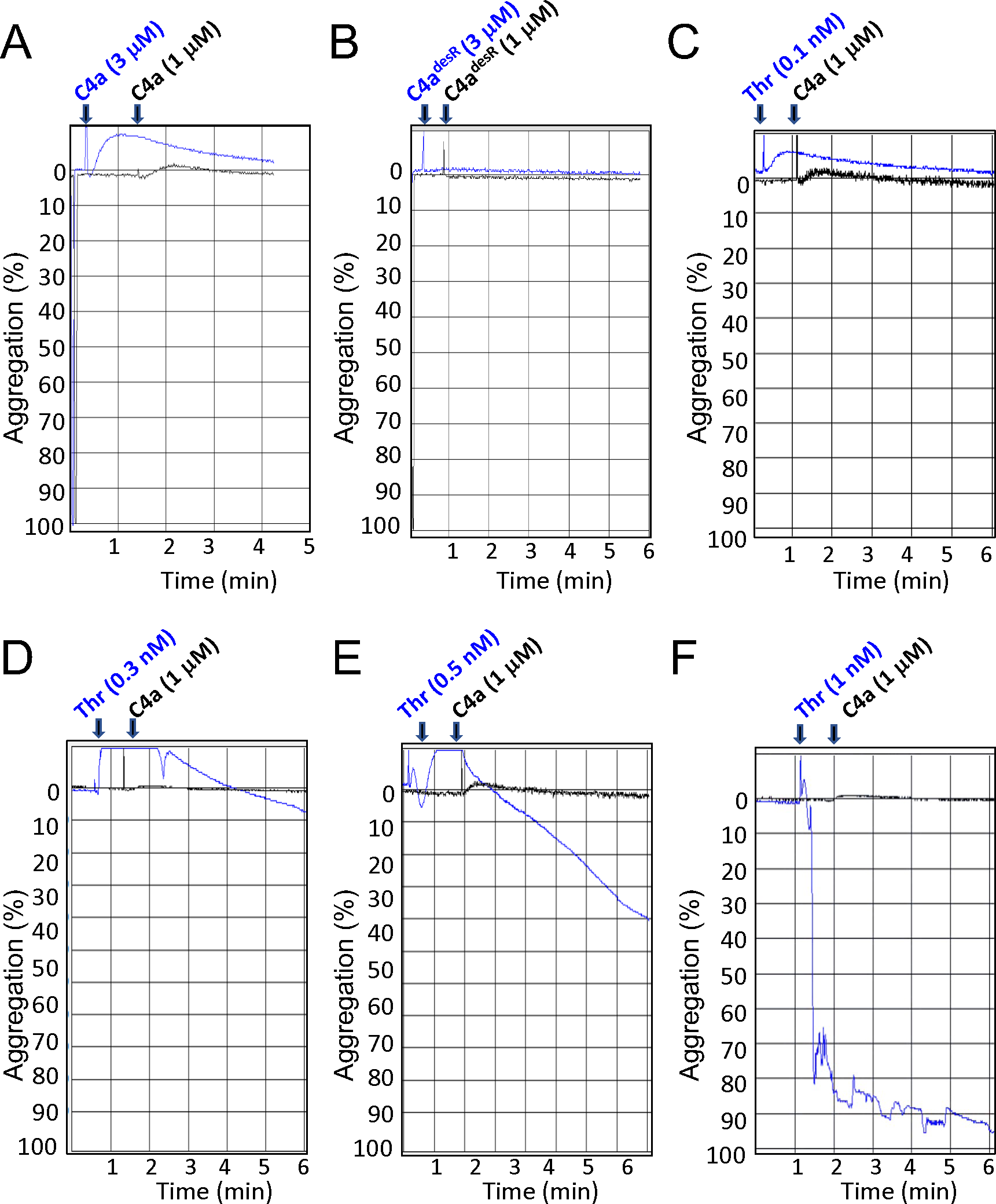
C4a and C4a^desArg^ fail to induce platelet aggregation. Human platelet rich plasma (PRP) and platelet poor plasma were prepared as described in materials and method section. The platelet aggregation was monitored by aggregometer (CHRONO-LOG® Model 700). **A**. C4a at the concentration of 1 or 3 μM fails to induce platelet aggregation. **B.** C4a^desArg^ at the concentration of 1 or 3 μM fails to induce platelet aggregation. **C**. Thrombin (0.1 nM) fails to induce platelet aggregation. **D**. Thrombin (0.3 nM) causes ∼10% platelet aggregation. **E.** Thrombin (0.5 nM) causes ∼ 40% platelet aggregation. **F.** Thrombin (1 nM) causes ∼ 100% platelet aggregation. The data is one representative from at least three independent experiments.

### C4a^desArg^ elicits [Ca^2+^]_i_ influx in HMEC-1 cells

Prior study showed C4a could trigger [Ca^2+^]_i_ influx in HMEC-1 cells ^(10)^. To determine whether C4a^desArg^ can elicit [Ca^2+^]_i_ influx in human HMEC-1 cells, Fluo-4 NW calcium assay kit was used to test [Ca^2+^]_i_ influx after C4a^desArg^ treatment. As shown in Fig. 2, similar to C4a, C4a^desArg^ could trigger [Ca^2+^]_i_ influx in a concentration-dependent fashion in human HMEC-1 ECs, but with decreased efficacy as compared to C4a ^(10)^. C4a^desArg^ can initiate [Ca^2+^]_i_ influx starting at the concentration as low as 10 nM.

**Figure 2.**
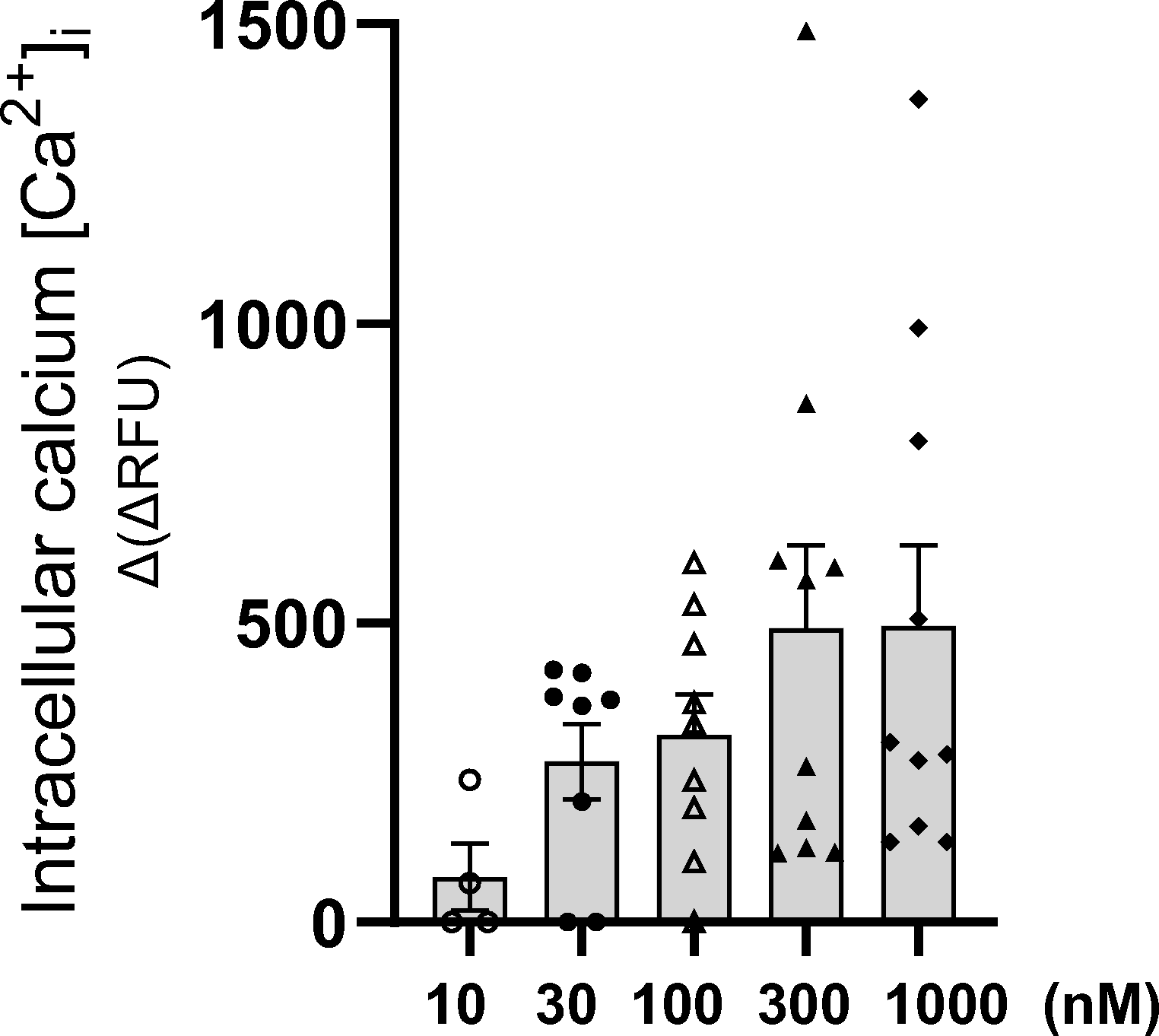
C4a^desArg^ induces [Ca^2+^]_i_ influx in HMEC-1 cells in a concentration-dependent fashion. The data are expressed as relative fluorescence [Δ(ΔRFU)] and represent the mean ± SE of four to seven independent experiments (n= 4-7 independent experiments).

### C4a^desArg^-induced ERK phosphorylation is PAR4-dependent

Prior study showed that C4a could induce ERK phosphorylation in a time- and concentration-dependent manner in HMEC-1 ECs. In addition, C4a-induced ERK phosphorylation is both PAR1- and PAR4-dependent as well as Gα_i_-independent ^(10)^. In the next experiment, we want to address whether C4a^desArg^ can cause ERK phosphorylation. As shown in Fig. 3A, C4a^desArg^ can significantly induce ERK activation (maximum ∼1.5-fold change of the control). The efficacy of C4a^desArg^-induced ERK activation is weaker than that of C4a (∼2.5-fold change of the control) ^(10)^, which is in line with the observation that the efficacy of C4a^desArg^-induced [Ca^2+^]_i_ influx (Fig. 2) and PAR1/4 activation in GPCR screening assay was weaker than that induced by C4a ^(10)^. Our results strongly suggest that the absence of arginine in the C-terminus of C4a will decrease its efficacy on [Ca^2+^]_i_ influx as well as ERK phosphorylation in HMEC-1 cells.

**Figure 3.**
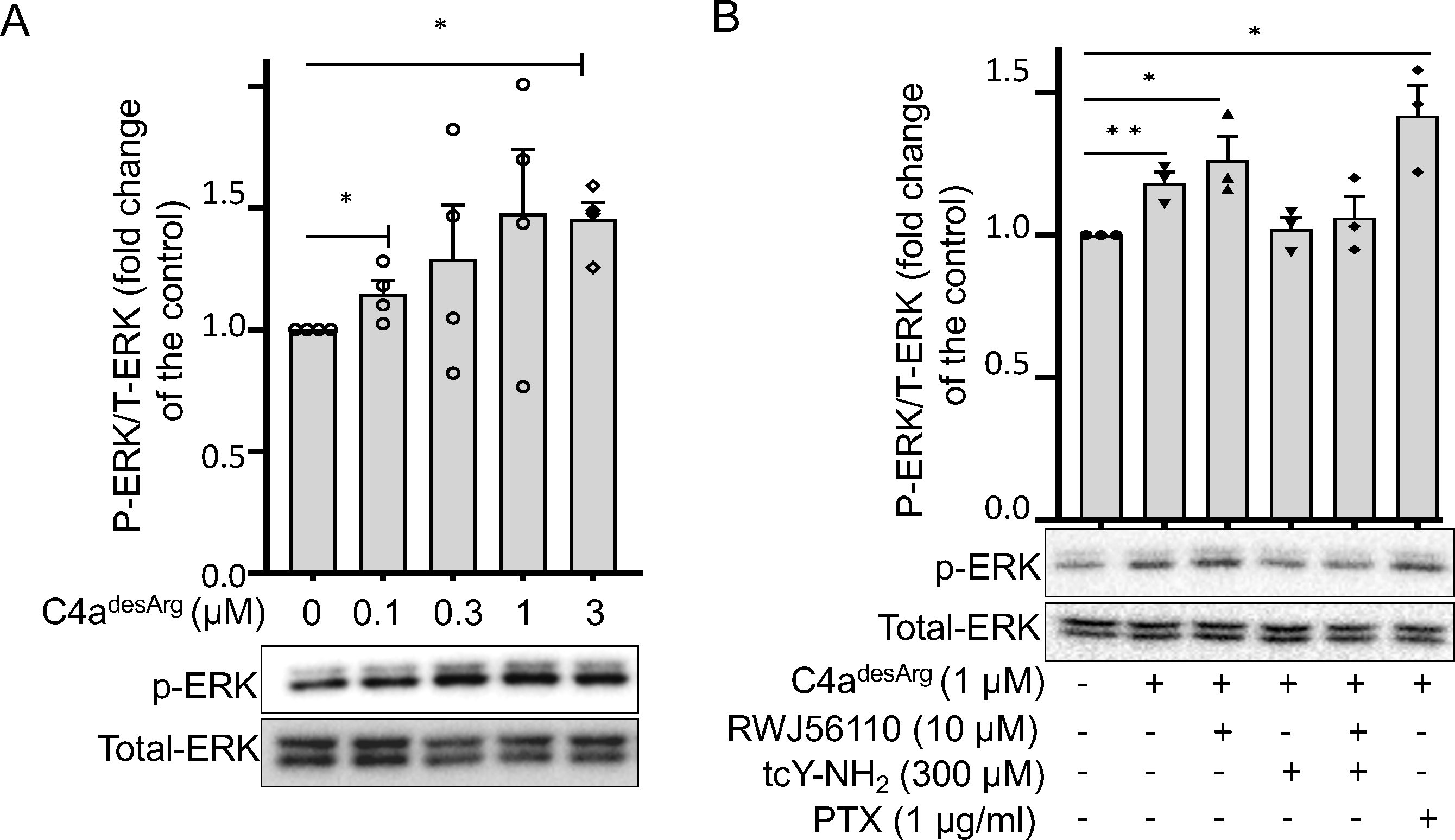
C4a^desArg^-induced ERK phosphorylation is PAR4-dependent and Gαi-independent signaling. **A**. C4a^desArg^ concentration-dependently enhances ERK phosphorylation. **B.** PAR4 antagonist tcY-NH2 (300 μM, 30 min), but not pertussis toxin (0.3 μg/ml, 10 h) and RWJ56110 (10 μM, 30 min) inhibit C4a^desArg^-induced ERK activation. The data are expressed as –fold change in densitometry of Western blots from control group and represent the mean ± SE of three independent experiments (n=3; \**p* < 0.05 *vs.* control; pairwise two-sided Student’s *t* test).

To determine the contribution of PAR1 and PAR4 on C4a^desArg^-mediated ERK phosphorylation, HMEC-1 cells were pretreated with respective antagonist for PAR1 (RWJ56110, 10 μM), PAR4 (tcY-NH_2_, 300 μM), or both antagonists for 30 min before adding C4a^desArg^. As shown in Fig. 3B, distinct from C4a, C4a^desArg^-induced ERK phosphorylation can be inhibited by PAR4 antagonist (tcY-NH_2_), but not by PAR1 antagonist (RWJ-56110). The inhibitory effect of both antagonists on ERK phosphorylation is like that of PAR4 antagonist alone (Fig. 3B). In addition, similar to C4a, C4a^desArg^-induced ERK phosphorylation is not affected by pre-treatment of Gα_i_ inhibitor pertussis toxin (PTX) in human HMEC-1 ECs (Fig. 3B), implying C4a^desArg^-induced ERK phosphorylation is also Gαi-independent ^(10)^. Our present study established that C4a^desArg^-induced ERK phosphorylation is PAR4-dependent and Gα_i_-independent. It seems that C-terminal arginine of C4a plays a role in determining the specificity to its receptors.

In summary, the data revealed that C4a^desArg^ acts similarly to C4a, which could trigger ERK phosphorylation and [Ca^2+^] influx in HMEC-1 ECs, but with the decreased efficacy as compared to C4a. In addition, different from C4a, C4a^desArg^-induced ERK phosphorylation is PAR4-dependent, indicating C-terminal arginine in C4a plays a role in governing the interaction of C4a with PAR1 and/or PAR4 and the downstream effector functions. Further structure-relationship of C4a^desArg^-PAR1/4 interaction needs to be characterized.

### C4a and C4a^desArg^ act differently from thrombin on Akt phosphorylation in human ECs

Thrombin is a typical activator for PAR1 and PAR4 by cleaving the N-termini of the receptors and the newly exposed tethered ligands bind to the receptors to cause downstream signaling effector functions, such as ERK phosphorylation and [Ca^2+^]_i_ influx in platelets and ECs^(19)^. Synthetic six-amino acid peptide mimic of PAR-1 or PAR-4’s N-terminal tethered ligand is sufficient to activate the receptor and induce similar cellular responses without proteolytic cleavage of the receptor ^(17)^. Our studies established that C4a and C4a^desArg^ act as untethered ligands for PAR1 and PAR4 and cause downstream ERK phosphorylation and [Ca^2+^]_i_ influx ^(10)^. It is still obscure whether thrombin and C4a/C4a^desArg^ share the same activation mode on PAR1 and/or PAR4 and cause the same downstream signaling effector functions. To address this question, we treated human ECs with thrombin at the concentration of 0.01, 0.03, 0.1, 0.3, and 1 nM to monitor the phosphorylation of ERK and Akt in Ea.hy 926 cells. As shown in Fig. 4A, thrombin at the concentration from 0.1 to 1 nM could concentration-dependently cause ERK phosphorylation (Fig. 4A *second row*), whereas Akt phosphorylation (Ser473) were significantly reduced as the concentration of thrombin increased (Fig. 4A *first row*).

**Figure 4.**
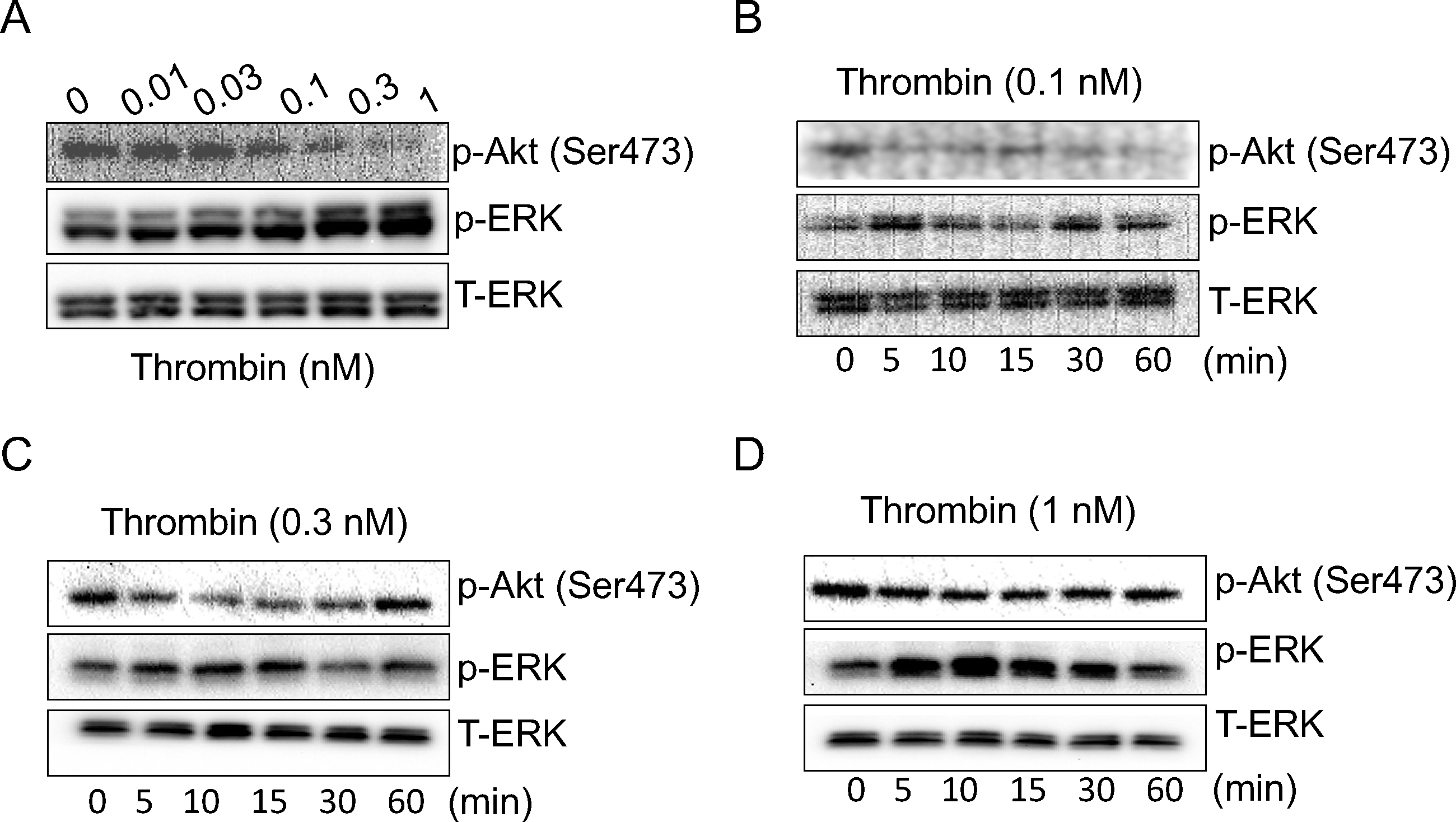
Thrombin causes ERK phosphorylation and Akt de-phosphorylation in human ECs. **A**. Thrombin concentration-dependently induces Akt de-phosphorylation and ERK phosphorylation. **B**. Time course of thrombin (0.1 nM) induces Akt de-phosphorylation and ERK phosphorylation. **C**. Time course of thrombin (0.3 nM) induces Akt de-phosphorylation and ERK phosphorylation. **D**. Time course of thrombin (1 nM) induces Akt de-phosphorylation and ERK phosphorylation. One representative Western blotting result from at least three independent experiments is shown. All experiments showed the same tendency.

In the next set of experiments, we tested time-course of thrombin’s effect at the respective concentration of 0.1, 0.3, and 1 nM on the phosphorylation of Akt and ERK. As shown in Fig. 4, Akt dephosphorylation (Ser473) starts from 5 min after the treatment of thrombin at the concentration of 0.1 nM (Fig. 4B), 0.3nM (Fig. 4C), and 1 nM (Fig. 4D) (*first row of each figure*). Conversely, ERK phosphorylation (*second row of each figure*) was significantly increased 5 min after the treatment of thrombin at the concentration of 0.1, 0.3, or 1 nM until 30 min.

In the next set of experiments, we tested the time-course of C4a (0.3 μM) and C4a^desArg^ (1 μM) on the phosphorylation of ERK and Akt in human ECs. As shown in Fig. 5, in contrast to thrombin Akt phosphorylation was significantly increased after the treatment of C4a (0.3 μM) (Fig. 5A) and of C4a^desArg^ (1 μM) (Fig. 5B) from 5 to 30 min in Ea.hy 926 cells (*first rows in* Fig. 5A and 5B). C4a/C4a^desArg^ can also cause ERK phosphorylation in human Ea.hy 926 cells (*second rows in* Fig. 5A and 5B). Similarly, Akt phosphorylation (Ser473) induced by C4a (0.3 μM) and C4a^desArg^ (1 μM) was also observed in a different human endothelial cells (HMEC-1) (Fig. 5C and 5D). In summary, the data shown that C4a/C4a^desArg^ act differently from thrombin in Akt phosphorylation. C4a and C4a^desArg^ induce significant Akt phosphorylation (Ser473), whereas thrombin causes significant Akt de-phosphorylation, indicating their discrete activation modalities on the receptor PAR1 and/or PAR4 and downstream signaling effector functions.

**Figure 5.**
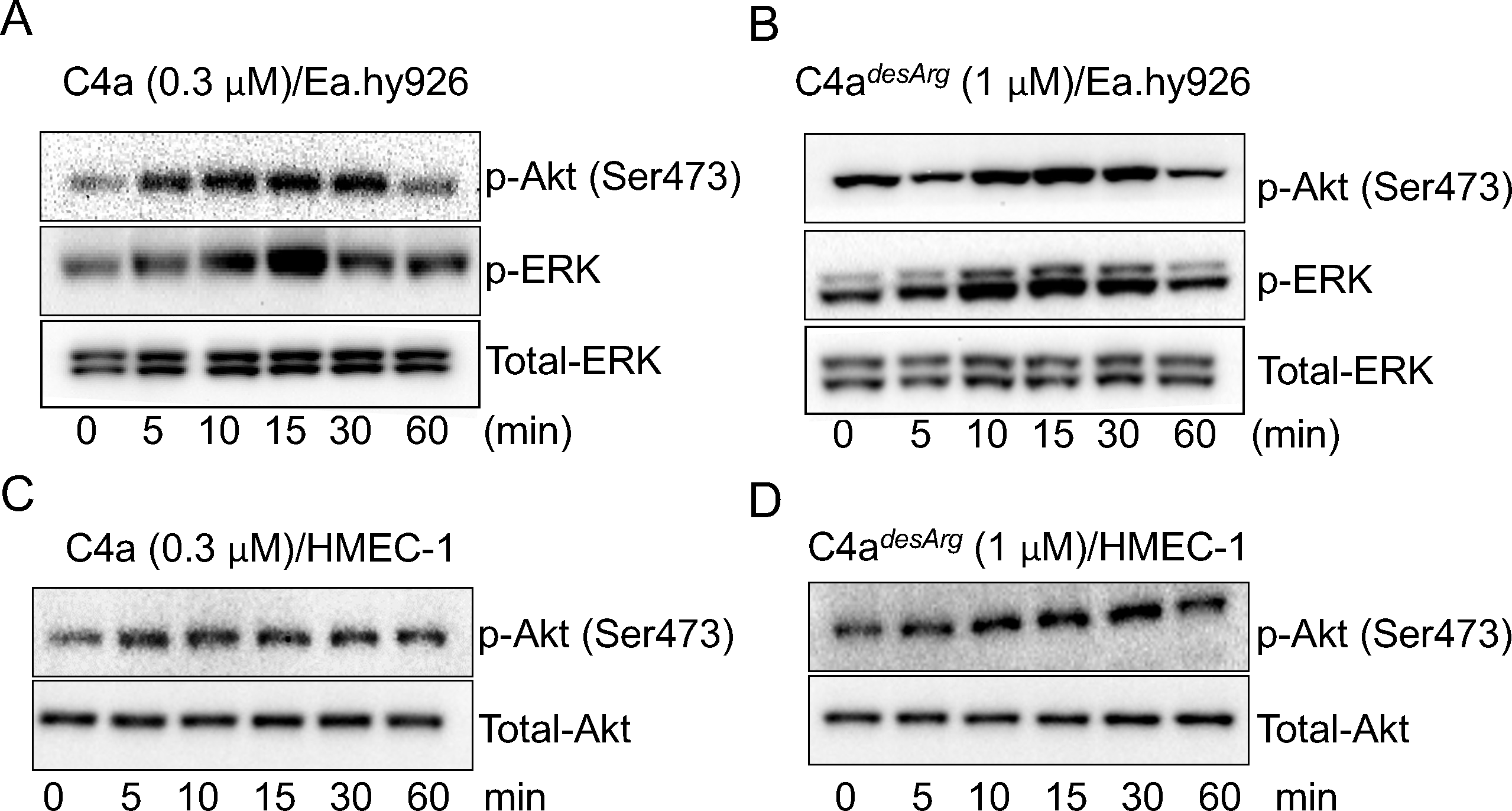
C4a and C4a^desArg^ induce phosphorylation of both Akt (Ser473) and ERK in HMEC-1 and Ea.hy926 cells. **A** and **C**. C4a (0.3 μM) triggers Akt phosphorylation in Ea.hy926 and HMEC-1 ECs. **B** and **D**. C4a^desArg^ (1 μM) triggers Akt phosphorylation in Ea.hy926 and HMEC-1 ECs. One representative Western blotting result from at least three experiments is shown. All experiments showed the same tendency.

### C4a^desArg^-induced Akt phosphorylation is PAR4-dependent

Our studies showed that C4a-induced ERK activation is PAR1 and PAR4 dependent ^(10)^, and C4a^desArg^-induced ERK activation in PAR4-dependent only (Fig. 3B). In the next experiment, we want to address whether C4a- and C4a^desArg^-induced Akt phosphorylation is PAR1 and/or PAR4 dependent. Human HMEC-1 cells were pretreated with PAR1 or PAR4 antagonist for 30 min and then the cells were treated with C4a (0.3 μM) or C4a^desArg^ (1 μM) for 10 min. As shown in Fig. 6 (*first row*), C4a-induced Akt phosphorylation could be inhibited by both PAR1 and PAR4 antagonists, whereas C4a^desArg^-induced Akt phosphorylation could be inhibited by PAR4 antagonist but not PAR-1 antagonist. We conclude that while C4a-induced Akt phosphorylation is both PAR1- and PAR4-dependent, C4a^desArg^-induced Akt phosphorylation is solely PAR4-dependent. The experiment recaptured the data that C4a-induced ERK activation is both PAR1- and PAR4-dependent, whereas C4a^desArg^-induced ERK phosphorylation is PAR4-dependent (Fig. 6, *second row*). Our data further established that C4a and C4a^desArg^ are the tethered ligands for PAR1 or/and PAR4 and C-terminal arginine is crucial in governing C4a signaling to PAR1 and/or PAR4 and downstream signaling effector functions.

**Figure 6.**
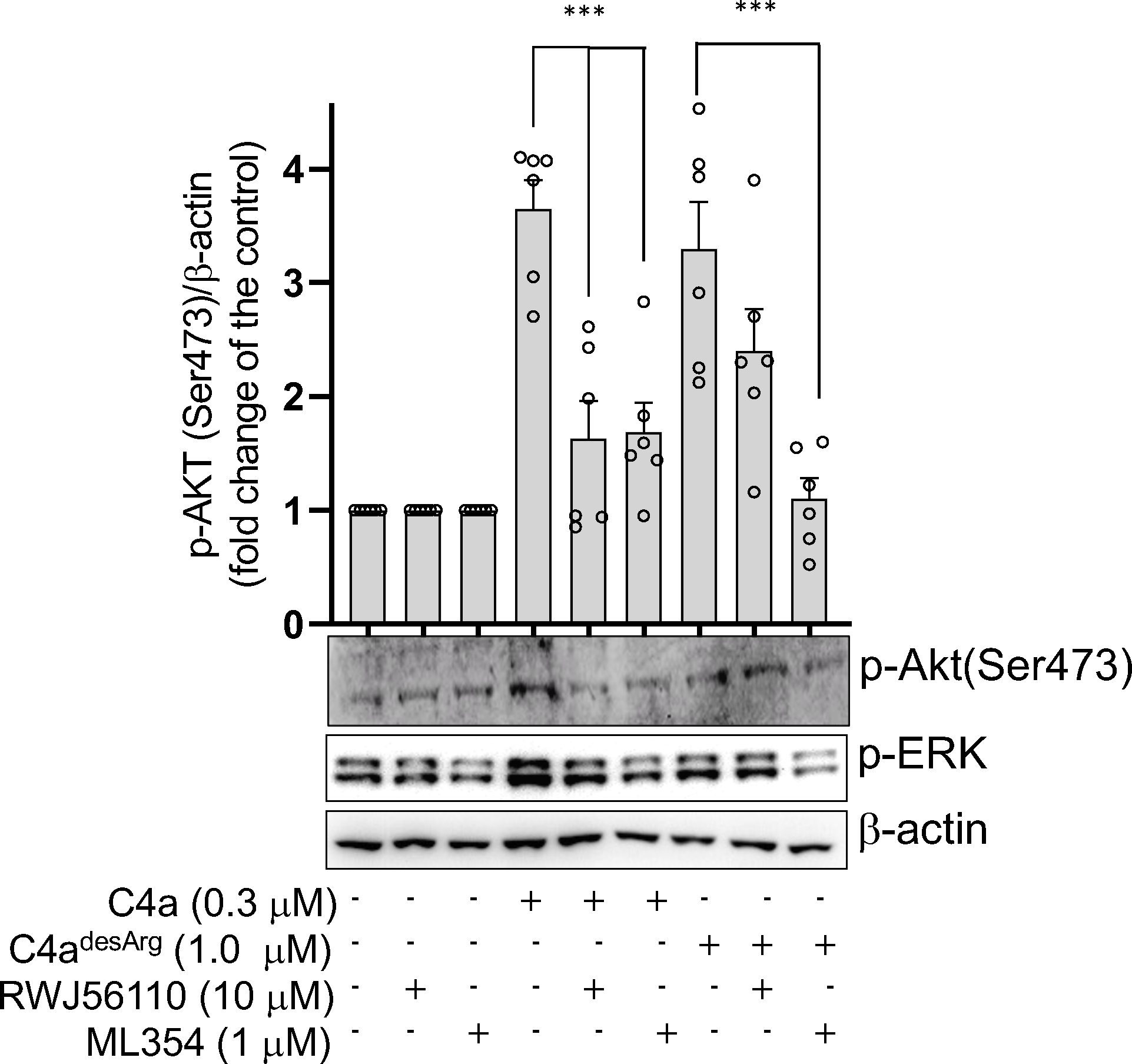
In HMEC-1 ECs, C4a-induced Akt phosphorylation (Ser473) can be inhibited by both PAR1 and PAR4 antagonists. However, C4a^desArg^-induced Akt phosphorylation (Ser473) can be inhibited by PAR4 antagonist but not PAR1 antagonist. **A**. Western blotting results show pretreatment of antagonists of PAR1 (*lane 5*) and PAR4 (lane 6) significantly inhibit C4a-induced Akt phosphorylation (Ser473) (*lane 4*). C4a^desArg^-induced Akt phosphorylation (Ser473) (*lane 7*) can be inhibited by the pretreatment of PAR4 antagonist (*lane 9*), but not inhibited by PAR1 antagonist (*lane 8*). One experiment from three separate experiments is shown. **B**. Densitometry analyses represent the mean ± SE of four independent experiments (n=4; \**p* < 0.05 *vs.* control; pairwise two-sided Student’s *t* test).

### Argatroban reverses thrombin induced Akt de-phosphorylation

Argatroban is a direct thrombin inhibitor that binds avidly and reversibly to the catalytic site of thrombin and that does not require other cofactors to exert its antithrombotic action ^(20)^. The thrombin induced Akt de-phosphorylation has been reported in human endothelial cells elsewhere ^(21, 22)^. In the experiment, we want to determine whether thrombin induced Akt de-phosphorylation is affected by Argatroban, a thrombin inhibitor. Our data clearly shown that Argatroban can reverse thrombin induced Akt de-phosphorylation at the concentration of 4 to 40 nM in human ECs (Fig. 7A). Our data showed that thrombin-induced Akt de-phosphorylation depends on its protease activity.

**Figure 7.**
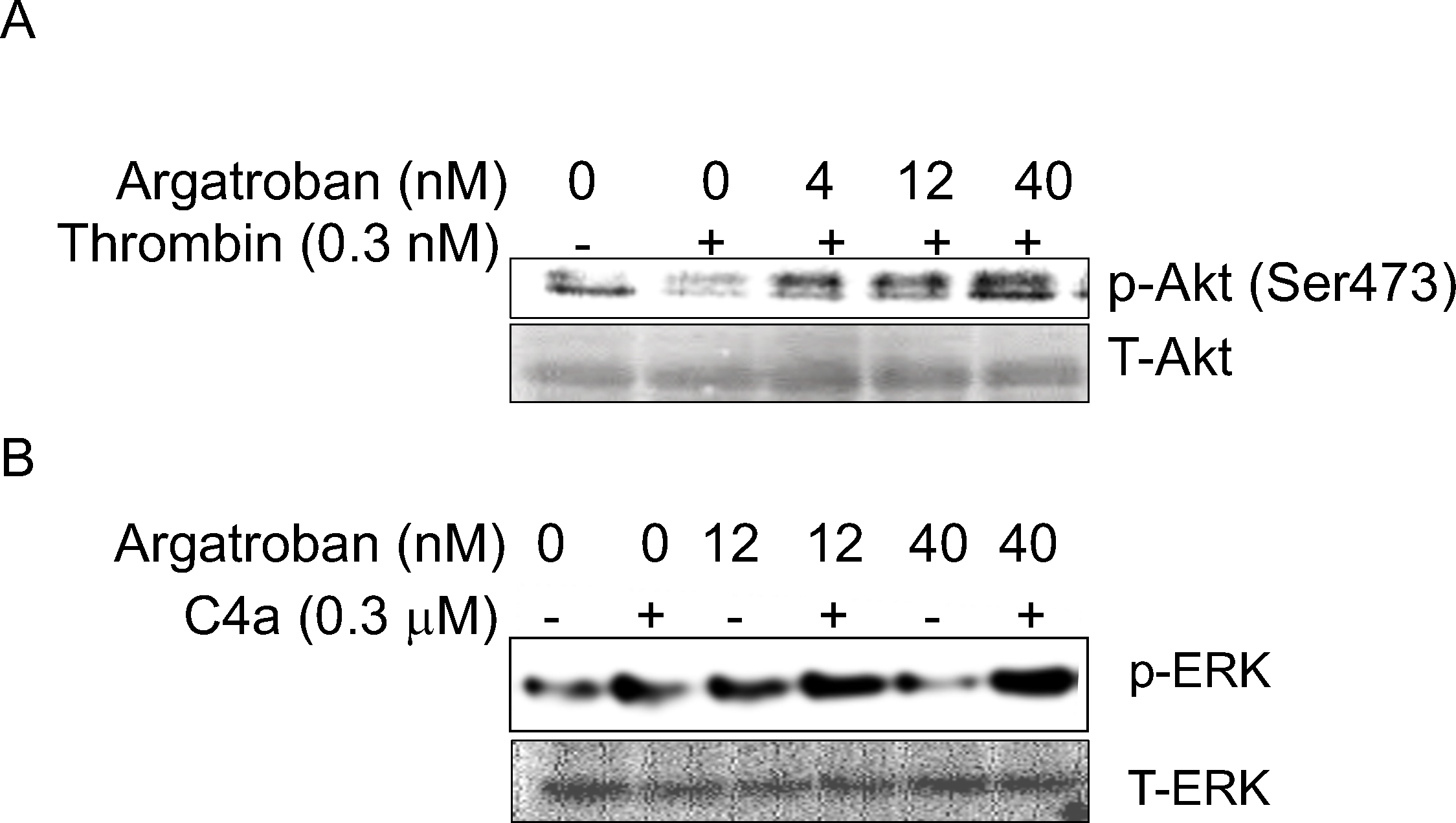
Thrombin inhibitor, Argatroban didn’t inhibit C4a-induced ERK phosphorylation at the concentration that can inhibit thrombin-induced Ak-de-phosphorylation in Ea.hy 926 human ECs. **A**. Argatroban (4, 12, and 40 nM) reverses thrombin-induced Akt de-phosphorylation. **B**. Argatroban (12 nM and 40 nM) has no effects on C4a-induced ERK phosphorylation. Each experiment has been done at least three times with the same tendency.

### Argatroban doesn’t affect C4a-induced ERK phosphorylation

As shown in Fig.1, C4a (3 μM) can trigger the similar effects as thrombin (0.1 nM) on platelet morphological change (form a negative “bump”). One speculation is that C4a-induced ERK phosphorylation could be due to the trace amount of thrombin in C4a preparation. To address this concern, we test whether Argatroban can reverse C4a-induced ERK phosphorylation. At first, we titrate the concentration of Argatroban at which it can reverse thrombin induced Akt de-phosphorylation. As shown in Fig. 7A, Argatroban can reverse the effect of thrombin-induced Akt de-phosphorylation at the concentration from 4-40 nM, which indicates that Argatroban at the concentration (4 to 40 nM) is enough to shut down 0.3 nM thrombin protease activity. In the next experiment, we want to test whether Argatroban at the concentration (4 to 40 nM) can inhibit C4a-indcued ERK activation. As shown in Fig. 7B, Argatroban did not affect C4a-induced ERK phosphorylation at the concentration of 12 nM and 40 nM, which could completely revers thrombin-induced Akt de-phosphorylation (Fig. 7A). We conclude that C4a-induced ERK activation through PAR1 and/or PAR4in human ECs was not due to the trace contamination of thrombin but the effect of C4a.

## Discussion

The activation of the classical or lectin complement pathway leads to the respective activation of C1s and MASP2, which can cleave the amino-terminal part of the α-chain at a single site of complement C4 to generate C4a (∼9 kDa) and C4b (∼195 kDa)^(2, 23)^. The arginine in the carboxyl terminal of C4a can be rapidly cleaved in the plasma by carboxypeptidases B and N to produce a stable des-arginated form of C4a (C4a^desArg^)^(11, 24)^. We posit that C4a^desArg^ is more relevant to the systemic functions/effects than C4a since C4a^desArg^ is more stable and can circulate. In contrast, C4a may produce the effects at the site where the classical or lectin complement pathway is activated. Our prior GPCR activation assay revealed C4a^desArg^ as a ligand for PAR1 and PAR4 but with less efficacy as compared to C4a. The present data clearly showed that C4a^desArg^ could exert effector signaling in human ECs at concentrations as low as ∼10 nM, which could trigger [Ca^2+^]_i_ influx and ERK phosphorylation. C4a concentration in plasma varies from ∼8 nM (healthy control) to ∼13 nM (patients with neovascular age-related macular degeneration) measured by a cytometric beads assay^(25)^. Another study using ELISA found the concentration of C4a in healthy controls, patients with chronic hepatitis B infection, and patients with liver failure related to chronic hepatitis B infection was 277 nM, 266 nM, and 209 nM respectively^(26)^. Our study revealed that C4a or C4^adesArg^ at the concentration of 300 nM almost produce the maximum signaling effector functions in human ECs, strongly indicating C4a and C4a^desArg^ might be responsible for some physiological and/or pathological functions. Recent studies revealed that C4a^desArg^ levels were significantly high in patients with IgA nephropathy and idiopathic pulmonary arterial hypertension (IPAH), which can serve as plasma protein biomarkers for complement activation in those disorders^(12, 13)^. It has been well established that the des-arginated forms of anaphylatoxins C3a and C5a (C3a^desArg^ and C5a^desArg^) usually either lack or have much less efficacy compared to their intact forms of anaphylatoxins^(24)^. We have demonstrated both C4a and C4a^desArg^ as untethered ligands for PAR1 and PAR4 to recruit β-arrestin in CHO-K1-PAR1 and CHO-K1-PAR4 engineered cell lines. As compared to C4a, C4a^desArg^ showed much less efficacy in those GPCRs activation assay^(10)^. In line with this observation, C4a^desArg^, like C4a, was observed to induce effector functions on ERK phosphorylation and [Ca^2+^]_i_ influx in human ECs but with significantly reduced efficacy as compared to C4a. The data indicated that the C-terminal arginine of C4a may play a role in governing its affinity and specificity to PAR1 and/or PAR4. Indeed, by applying the respective antagonist of PAR1 and PAR4 before C4a^desArg^ treatment, our data verified that C4a^desArg^-induced ERK phosphorylation can only be inhibited by the PAR4 antagonist, but not by the PAR1 antagonist, which is distinctive from C4a that can induce ERK phosphorylation through both PAR1 and PAR4. The data suggest that the C-terminal arginine may govern the interaction of C4a with PAR1 and/or PAR4. The role of arginine in the interaction of C4a with PAR1 and PAR4 needs to be elucidated in future studies.

Once establishing C4a as a ligand for PAR1 and PAR4, we immediately initiated to investigate whether C4a and C4a^desArg^ could activate PAR1 or PAR4 to trigger platelet aggregation. Unexpectedly, we found that C4a and C4a^desArg^ were unable to trigger platelet aggregation even at supraphysiological concentrations (3 μM) (Fig. 1A and 1B), which was supported by a recent study by Han *et al.*^(18)^. It is still unclear why C4a and C4a^desArg^ were unable to trigger platelet aggregation although they can bind to PAR1 and PAR4 as an untethered ligands and trigger effector functions in human ECs^(10)^. Han *et al.* proposed that C4a mediates selective activation of PAR1 and/or PAR4, which might involve specific membrane localization, cofactors, or both ^(18)^.

Defining a specific receptor for C4a has long been a challenge and obscure. Prior studies revealing its low affinity to other anaphylatoxin receptors raise the question of whether C4a is involved in physiological significance^(3, 8, 9)^. Prior GPCR activation assays screening for receptors for C4a using reporters with 168 known and 73 orphan GPCRs have changed the game, from which PAR1 and PAR4 were identified as receptors for C4a as well as C4a^desArg(10)^. Thrombin has been well established as a typical ligand for PAR1 and PAR4. Thus, the exclusion of possible trace thrombin contamination in the preparation of C4a is very essential. Our prior study applied a chromogenic substrate cleavage assay (Sensolyte 520 Thrombin activity assay kit) to check the possibility of thrombin contamination in C4a preparation (Complement Tech. Taylor, Texas). We couldn’t detect thrombin activity in C4a preparation even at a concentration of 3 μM using the chromogenic substrate cleavage assay. Nonetheless, the detection limit of this kit is around 0.1 nM of thrombin. In addition, our present platelet aggregation study revealed that platelet morphological change (a negative “bump”) triggered by C4a (3 μM) resembles the effects induced by thrombin (0.1 nM) (Fig. 1A and 1C). These observations lead us to speculate possible trace contamination of thrombin in C4a preparation, which might generate all signaling effector functions, such as ERK activation and [Ca^2+^]i influx ^(10)^. To address this issue, we titrated the effect of thrombin with a series of concentrations (0.01, 0.03, 0.1, 0.3, and 1 nM) on the phosphorylation of ERK and Akt in human ECs. We found that thrombin at the low concentrations (0.01 and 0.03 nM) caused no significant effects on ERK phosphorylation. At the higher concentrations of 0.1, 0.3, and 1 nM, thrombin can significantly induce ERK phosphorylation in a concentration and time-dependent manner; however, under the same conditions, it can significantly cause dephosphorylation of Akt (Fig.4). In parallel, we tested the effects of C4a and C4a^desArg^ on the phosphorylation patterns of Akt and ERK. Our data revealed that C4a (300 nM) and C4a^desArg^ (1 μM) could trigger the phosphorylation of both ERK and Akt (Fig.5), which showed a completely different downstream activation modality compared to thrombin. The data conclude that C4a and C4a^desArg^-induced effector functions aren’t caused by the possible trace contamination of thrombin. To further consolidate this result, we applied the thrombin-specific inhibitor, Agatroban to test if it can inhibit thrombin-induced Akt dephosphorylation but could not inhibit C4a-induced ERK phosphorylation. As shown in Fig.7, Agatroban at concentrations of 4 to 40 nM could significantly reverse thrombin (0.3 nM)-induced Akt dephosphorylation (Fig. 7A) but could not affect C4a-induced ERK phosphorylation (Fig. 7B), clearly demonstrating that no thrombin is involved in C4a-induced ERK activation.

Using respective antagonists for PAR1 and PAR4, a prior study demonstrated that C4a-induced ERK phosphorylation is PAR1- and PAR4-dependent and Gαi-independent in human ECs. Interestingly, the present study demonstrated that C4a^desArg^-induced ERK and Akt phosphorylation is PAR4 dependent in human ECs. PAR4 was primarily studied in the context of platelet functions, and it appears that PAR4 and PAR1 signaling pathways are redundant. Emerging evidence has shown that PAR4 has its unique activation pathway and plays a critical role in multiple patho- and physiological processes, such as platelet activation and aggregation, inflammation, and pain^(27–31)^. Upon receptor activation, while PAR1 can be phosphorylated on key serine residues causing its rapid internalization and degradation, PAR4 doesn’t go through the processes of phosphorylation, internalization, and degradation, which will prolong stimulus of the receptor. PAR4 can form a heterodimer with PAR1 and P_2_Y_12_ receptors to alter the signaling in platelets^(32)^. Early studies have established that C4a possesses a strong chemotaxis inhibitory effect on monocytes at concentrations as low as 10^−16^ M^(33)^. C4a was also reported to inhibit C3a-induced O_2_^·−^ generation in guinea pig macrophages. In addition, recombinant human C4a was recently demonstrated to impair C5a-induced neointima formation, reduce C3a- or C5a-mediated chemoattractant and secretagogue activities in mast cells, and prevent hyperoxic lung injury via a macrophage-dependent signaling pathway^(5–7)^. It appears that C4a/C4a^desArg^ may perform anti-inflammatory effects. One interesting paradigm could be that C4a/C4a^desArg^ generated from earlier component C4 in classical or lectin pathways might be able to alleviate the inflammatory effects caused by anaphylatoxins C3a and C5a generated from the activation of later complement components. Our future studies will explore the effects of C4a/C4a^desArg^-PAR1/PAR4 signaling pathway on the functions of monocytes, T, and B lymphocytes, which might play important roles in autoimmune and other disorders.

## Author Contributions

M.L. and H.W. designed experiments. M.L. V.L. and H.W. carried out the experiments. All authors agreed to the version of the manuscript.

## Acknowledgement

We appreciated Dr. John D. Lambris, Department of Pathology and Laboratory Medicine at the University of Pennsylvania Perlman School of Medicine for providing us reagents and equipment to carry out platelet aggregation assay.

